# Macro-Equi-Diff (MED): Scaffold-based Macrocycles Generation Using Equivariant Diffusion

**DOI:** 10.64898/2026.02.05.703948

**Authors:** Sai Shobit Kambampati, Siri Anumandla, Sai Lahari Guttula, Varshith Reddy Kavadi, Sanjana Gogte, Vani Kondaparthi

## Abstract

Macrocyclic compounds are essential in drug discovery as they can modulate protein–protein interactions and enhance selectivity. Their structural complexity enables access to molecular diversity beyond traditional small molecules; however, designing feasible macrocycles remains a challenging task. Current computational methods often fail to generate macrocycles with proper drug-like properties. Here, we present Macro-Equi-Diff (MED), a deep learning framework that combines transformer-based site identification with an E(3)-equivariant Diffusion Model (EDM) for linker creation, and a fragment–linker attachment module. MED transforms acyclic molecules into structurally consistent macrocycles. MED was tested on the ZINC dataset, achieving high validity (93.92%), uniqueness (99.94%), macrocyclization (99.92%), and linker novelty (82.81%). MED improves upon previous methods that lack a macrocyclic geometry context. As a case study, MED was used to macrocyclize four acyclic drugs targeting the JAK2 protein. The generated macrocycles exhibited favourable molecular descriptors and strong binding affinities, establishing MED as a reliable method for expanding the macrocyclic chemical space.

## 1. Introduction

Macrocycles, defined as cyclic molecules with rings of 11 or more atoms, occupy a distinctive space between small molecules and large biologics^1^. Their large ring size imparts conformational flexibility, enabling diverse three-dimensional shapes that can engage binding surfaces inaccessible to traditional small molecules^2^. This makes them particularly effective for targeting complex protein-protein interactions (PPIs)^3^, where small molecules lack sufficient affinity and biologics are often too large.

In drug discovery, macrocycles combine high affinity and selectivity with favorable pharmacological properties^4^. Their pre-organized conformations reduce the entropic penalty of binding. At the same time, their ability to shield polar groups is often described as the “chameleon effect,” which enhances permeability and oral bioavailability, even beyond the traditional Rule of Five (bRo5)^5,6^. Although many approved macrocycles are natural products, advances in synthesis are expanding this chemical space, enabling the design of novel scaffolds for next-generation therapeutics^7^.

Computational methods were employed to expedite the discovery of macrocyclic compounds. Classical approaches, such as rule-based algorithms and library-guided linker scans, ensured chemical validity but were compromised by a deficiency in structural diversity^8^. The application of artificial intelligence, specifically deep learning, has shown immense potential in various stages of the drug discovery pipeline^9^, including de novo molecular design, scaffold hopping, structural optimization, and activity prediction. Subsequent machine-learning strategies, such as sequence-to-sequence models, have attempted to “translate” acyclic structures into their corresponding macrocyclic forms by adding linkers^10^. However, because the process is handled only as a sequence translation task rather than as a chemically informed generative approach, these methods frequently fail in providing sufficient chemical context during training, which restricts the model’s capacity to capture underlying molecular interactions. They also struggle to learn proper SMILES^11,12^ syntax due to a lack of chemical context.

The transformer-based Macformer^10^ often lacks an explicit chemical context and treats macrocycle generation from acyclic precursors as a straightforward translation task. Additionally, unlike diffusion models, which can produce diverse outputs from the same input through stochastic sampling, MacFormer is deterministic and always yields identical results for the same inputs. Database-driven strategies, such as MacLS^13^, ensure validity but are constrained by the library content, which limits chemical diversity.

The MED framework is developed to address the above challenges. Unlike 2D SMILE sequence-to-sequence models, MED employs an EDM^14^ for the end-to-end design of linkers in 3D space, thereby enforcing both geometric and chemical plausibility. Instead of tapping predefined libraries, it generates brand-new linkers de novo^15^, thereby maximizing structural diversity. This enables MED to produce unique, stable, drug-like macrocycles, thereby extending computational means for targeting previously untreatable proteins.

A case analysis of clinically effective kinase inhibitors, including Fedratinib^16^, Ruxolitinib^17^, Sunitinib^18,19^, and Lenvatinib^20,21^, was conducted. Macrocycle analogues generated by the MED framework for these inhibitors displayed favorable docking scores against the JAK2 protein^22^ compared to their corresponding acyclic structures, indicating that their binding interactions could be enhanced through macrocyclization. These findings highlight the potential of MED in developing drug-like macrocycles with therapeutic promise.

## 2. Methodology

The macrocyclization framework is structured as a three-step computational pipeline for producing and predicting novel macrocycles, as shown in **Fig. 1**. The macrocyclization workflow in the MED model is given as an algorithm in the supplementary file.

**Fig. 1.**
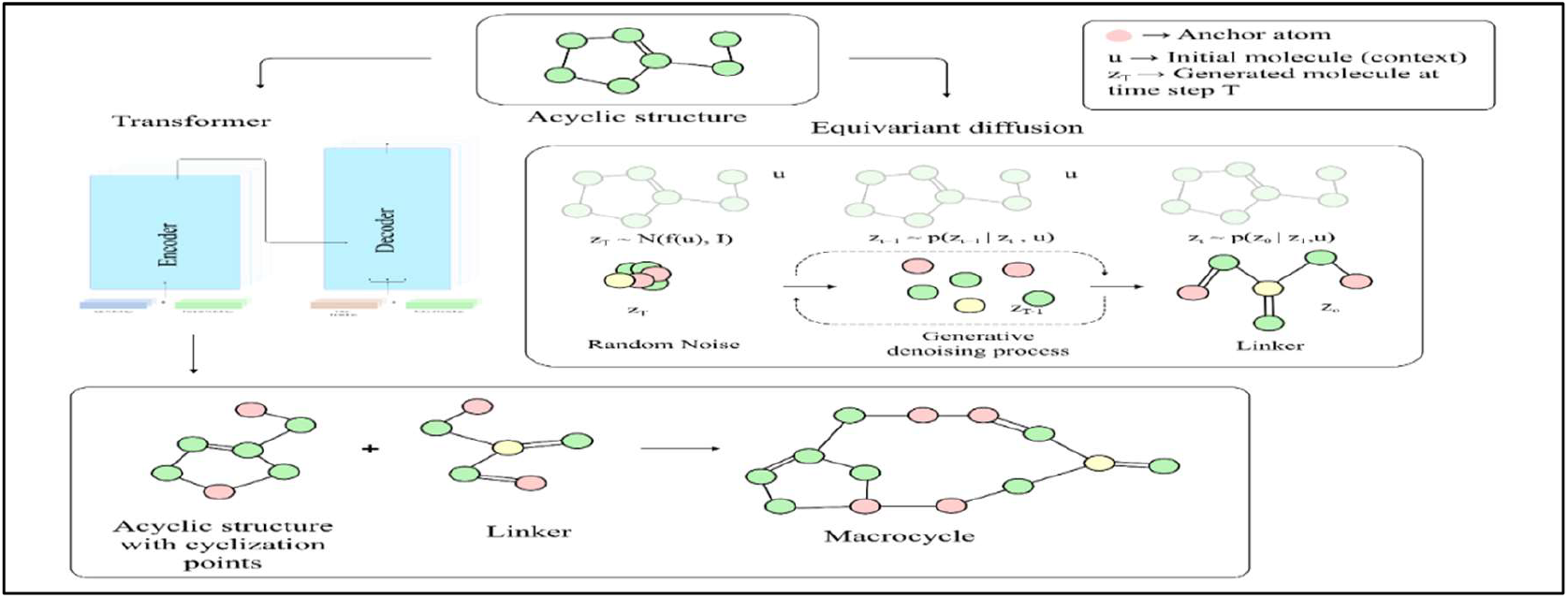
An overview of the MED macrocyclization framework. Figure 1 presents the process beginning with transformer-based site identification, in which two chemically valid attachment points are predicted on an acyclic molecule (highlighted in pink). These serve as anchor sites for macrocyclization (as represented in the linker). In the second stage, an EDM with an EGNN generates linkers conditionally on the acyclic molecule (described in the figure as u), ensuring geometric consistency and invariance to rotations/translations. The linker is then attached to the predicted anchor sites, yielding the final macrocycle. This three-stage pipeline enables scaffold-conditioned macrocycle design in a symmetry-preserving manner.

1. Transformer-based site identification for appropriate linker attachment points on acyclic SMILES.
2. EDM-based linker synthesis - a conditional generator model - creates linkers in an attempt to complete the macrocycle.
3. Prospective macrocycles are formed by connecting fragments and linkers.

### 2.1. Dataset and Training Setup

#### 2.1.1. Transformer Model Dataset

The dataset was preprocessed using the ChEMBL database^23^ and filtered to get valid macrocyclic structures. Each macrocycle was fragmented using a fragmentation script that breaks two single bonds in the largest ring to generate an acyclic fragment and its linker. The linker is constrained to 3–9 structural atoms, with no large rings (≥7 atoms), and ≤25% of the macrocycle’s structural atoms. Canonicalized SMILES of the acyclic fragments were used to build the dataset for the Transformer model. The input is in acyclic SMILES format without cyclization points, and the output includes dummy atoms (represented by *) that mark true cyclization sites. Since the acyclic structures are fragments of prevailing macrocycles, the model learns chemically valid and optimal placement of cyclization sites.

#### 2.1.2. Diffusion Model Dataset

For training the diffusion model, the dataset hosted at Zenodo (https://zenodo.org/records/7121278) is used. This is derived from the Geometric Ensemble Of Molecules (GEOM) dataset^24^. The fragment-linker pair dataset was divided into 282,602 pairs for training and 1,251 pairs for validation. EDM was then tested on 5,551 pairs from the ZINC dataset (https://github.com/yydiao1025/Macformer/tree/main/data/ZINC).

RDKit was used to create three-dimensional atomic coordinates from SMILES representations. The ETKDG algorithm was used to embed an initial conformer ensemble for each molecule and fragment. The Merck Molecular Force Field (MMFF) was then applied to minimize energy, and the lowest-energy conformer was kept as the representative three-dimensional structure. The diffusion model was fed these optimized geometries. In order to infer covalent bonds from interatomic distances, saturate open valences with hydrogens, and create standardized SDF files, OpenBabel was used to process the model’s raw atomic point clouds (.xyz format, encoding atom types and Cartesian coordinates without explicit bond connectivity). The same OpenBabel pipeline was used to process ground-truth reference structures in order to guarantee that downstream 3D metrics like RMSD represent real structural differences rather than file format artifacts.

### 2.2. Transformer-Based Attachment Point Prediction

A Transformer Encoder–Decoder architecture is used to accept SMILES of acyclic molecules and predict two chemically valid macrocyclization anchor points, which are marked with an asterisk (*). **Fig. 2** shows the overall model architecture and the transformer-based prediction of attachment sites.

**Fig. 2.**
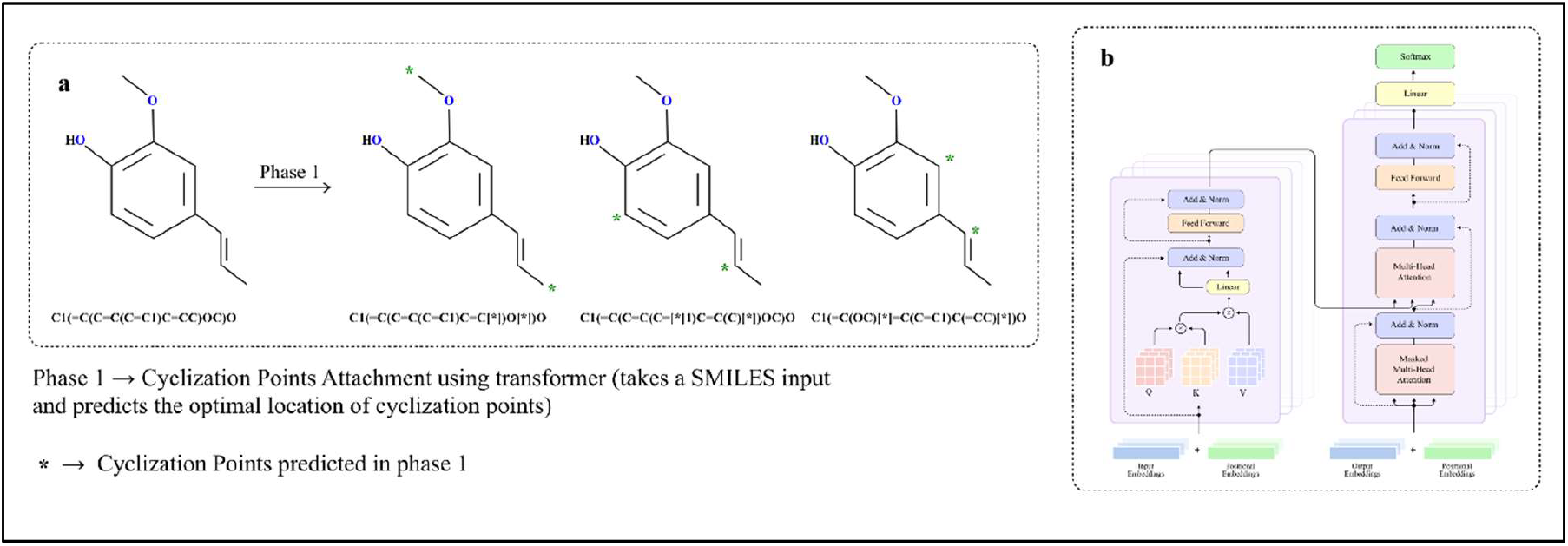
Transformer-based prediction of attachment sites and model architecture overview. **a**. Transformer-based attachment site prediction. **b**. Transformer architecture - A scaled-dot attention mechanism is given inputs in the form of three matrices: matrix Q containing a collection of queries, matrix K containing the keys, and matrix V containing the values, as explained in Figure 2.

#### 2.2.1. Transformer Model Architecture

The Transformer consists of an autoregressive encoder and decoder. Both input and output SMILES sequences are tokenized and transformed into a learnable embedding matrix, where a 256-dimensional vector represents each token. To encode sequence order, sine and cosine functions serve as positional encodings, with “pos” indicating the token’s position and “i” as the iterator from 0 to d_emb_/2 as shown in equation 1. The positional encodings are then added to the token embeddings, enabling the model to incorporate both positional information and semantic meaning.

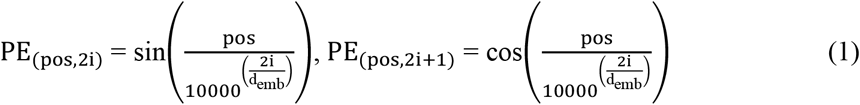

#### 2.2.2. Transformer Encoder and Decoder

The encoder processes the input SMILES sequence, extracting contextual information across tokens. Simultaneously, the decoder generates the target sequence by attending to both the encoder’s outputs and previously generated tokens. To preserve the autoregressive property, the decoder employs masked self-attention^25^. The architecture consists of four encoder layers, four decoder layers, four attention heads, a feedforward network size of 512, and a dropout rate of 0.1.

The multi-head attention mechanism^26^ enables the encoder and decoder to simultaneously focus on multiple tokens, allowing the Transformer to manage long-range dependencies efficiently. Each multi-head attention unit consists of 8 parallel scaled dot-product attention layers, which are concatenated and linearly projected to produce the final output. Each scaled dot-product attention layer receives three matrices as input: Q (queries), K (keys), and V (values). The attention is then computed as shown in Equation 2.

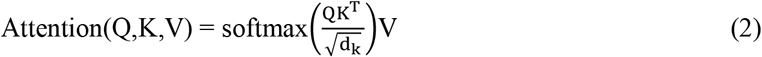

where d_k_ is a scaling factor based on the size of the weight matrices.

Finally, the Transformer’s output is passed through a fully connected layer that projects it onto the target vocabulary space, enabling token-level predictions. During inference, the model predicts one token at a time until the end-of-sequence token is generated or the maximum sequence length is reached.

Detailed training configurations and optimization hyper-parameters for the Transformer model are provided in the Supporting Information (Section S1.1).

Performance metrics for the model are reported in the Results and Discussion section.

### 2.3. Diffusion-Based Linker Generation using EGNN

The core of the macrocyclization framework is a diffusion generative model^27^ built with Equivariant Graph Neural Networks (EGNNs)^28,29^. The input to this module is the molecular graph of the acyclic molecule, along with constraints on the number of atoms in the linker.

#### 2.3.1. Diffusion Models Overview

Diffusion models are generative models^30^ that consist of two key processes:

##### Noising

A gradual noising of a data point x into a noise vector z_t_ over time steps t = 0, …, T as presented in **a** of **Fig. 3**. The equation is illustrated in Equation 3.

**Fig. 3:**
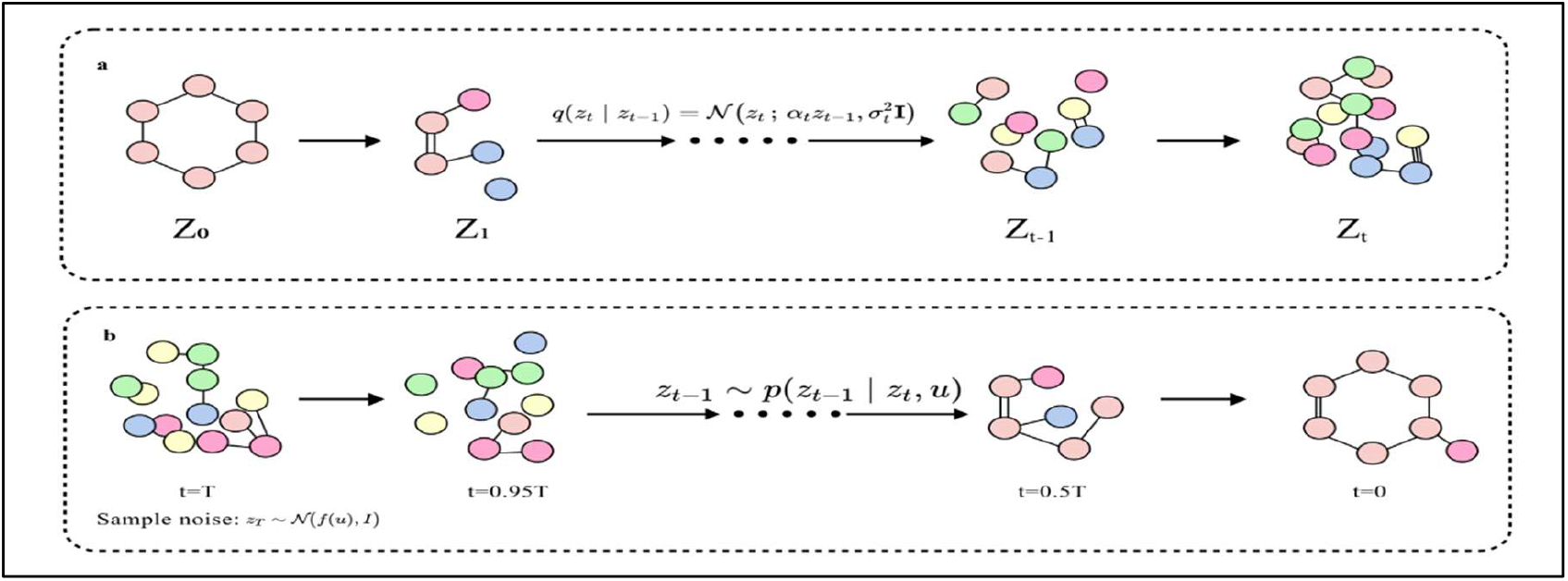
Applying the diffusion process to a molecular graph. **a**. Noising: The clean molecule z_O_ undergoes progressive noising through a Markov chain, where Gaussian perturbations are added at each step. As the time step t increases, both atomic positions and bond connectivity become increasingly corrupted, transforming the ordered structure into a highly noisy representation z_t_. Colored nodes denote atoms, and edges denote chemical bonds. **b**. Denoising: Starting from random noise, the diffusion model iteratively removes noise and reconstructs structural information by gaining context from the conditional input (u). This gradually recovers atomic positions and bond connectivity. The process ultimately yields a chemically valid molecular graph from the coordinate space.

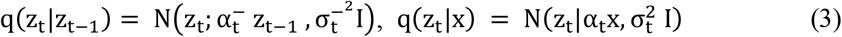

Here, 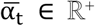 determines the extent of signal preservation, while 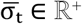 controls the amount of noise introduced. By hypothesis, the transition model follows the Markov property, as expressed in the equation 4.

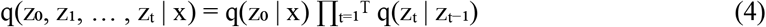

**Denoising:** A learned neural process that approximates the inversion of noise to reconstruct valid data from z_t_ as depicted in **b** of **Fig. 3**, is expressed in Equation 5.

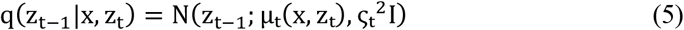

where,

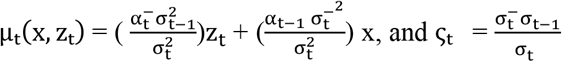

The second component of a diffusion model is the generative denoising process, which is trained to reverse the noising process when a data point x is not obtained. The generative transition distribution is formulated in Equation 6..

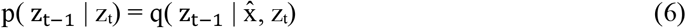

The data point x represents a molecular linker in coordinate space, and the model learns to reverse the noisy trajectory conditioned on the acyclic core molecule. Since the true x is unknown during generation, a neural network φ predicts the added noise 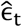, reconstructing x as:

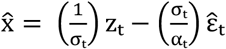The network minimizes mean squared error, 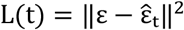with timesteps uniformly sampled.

#### 2.3.2. E(3)-Equivariant Graph Neural Network for Denoising

The learnable function φ, which models the dynamics of the reverse diffusion process, is implemented as a modified EGNN ^24^. A conditional distribution p(x∣y) is E(3)-equivariant if p(Rx+ t ∣ Ry + t) = p(x∣y) where R ∈ O(3) denotes a rotation or reflection and t ∈ R^3^ a translation. Similarly, for functions f: R^3^→R^3^, O(3)-equivariance requires f(Rx) = Rf(x), while O(3)-invariance of conditional distributions requires p(Rx ∣ Ry) = p(x∣y), where R denotes rotations or reflections and t translations. Translation invariance further requires that f(x + t) = f(x). As illustrated in **Fig. 4**, the network preserves equivariance under rotations and reflections (O(3) equivariance), enabling consistent reverse diffusion to generate chemically valid linker structures^31^.

**Fig. 4:**
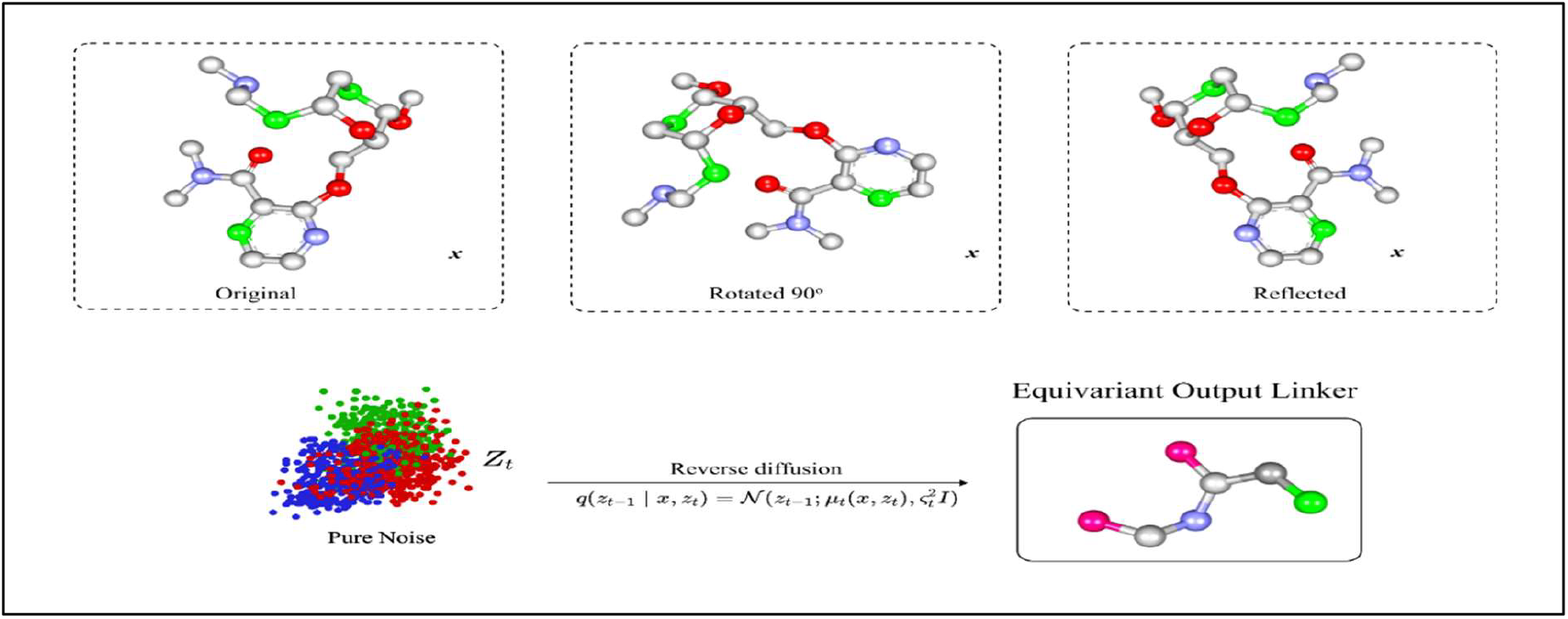
Illustration of O(3)-equivariance in the diffusion-based linker generation process. Top row: The same input scaffold in its original, rotated 90°, and reflected forms. Bottom row: The equivariant diffusion model denoises noisy inputs. While sampling is stochastic, outputs remain consistent up to the same rigid transformation. Figure 4 illustrates the O(3) component of equivariance, showing that rotations and reflections of inputs yield correspondingly transformed outputs, preserving geometric consistency. The final output is a linker structure ready to be connected to the scaffold, yielding a complete macrocyclic molecule.

The network explicitly incorporates the Center-of-Mass (COM) information from the context point u. For each input fragment, the COM of the anchors is computed and used to center the coordinates before applying φ, ensuring translation invariance. To achieve this, following Hoogeboom et al. (2022), the initial coordinates are subtracted from the predicted coordinate noise as depicted in Equation 7.

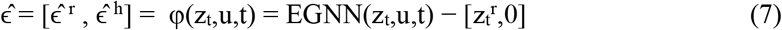

This ensures that the predicted noise is computed relative to the COM, effectively removing any dependence on the absolute positions of the molecule or its fragments.

These two components are merged into a single fully connected molecular graph:

- Nodes are represented by their coordinates r_i_∈R^3^ and feature vectors h_i_ that encode atom type, time step t, and fragment flags.
- The network predicts a noise vector 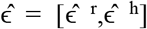 with both coordinate and feature components.

As diffusion sampling is stochastic, individual outputs may differ in atomic arrangement, but their distributions remain related by the same geometric transforms applied to the inputs.

#### 2.3.3. EGNN Layer Formulation

The EGNN is composed of Equivariant Graph Convolutional Layers (EGCLs)^32^, each of which updates coordinates and features jointly, as formulated in Equation 8.

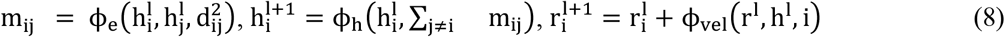

Where, 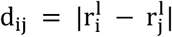 is the interatomic distance, ϕ_e_ and ϕ_h_ are learnable functions (MLPs)operating on invariant features, ϕ_vel_ updates coordinates in an equivariant manner. Equivariance is preserved by construction:

- ϕ_e_ and ϕ_h_ depend only on E(3)-invariant quantities (node features, pairwise distances).
- ϕ_vel_ depends linearly on coordinate differences (r_i_ − r_j_), ensuring E(3)-equivariance.

After applying a sequence of EGCLs, the network produces updated coordinates and features,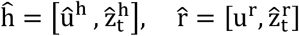 The context-related outputs, and only 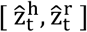, are retained as the EGNN’s final output.

The average Tanimoto similarity between the generated linkers and the linkers in the diffusion model training set is shown in Supplementary Fig. SF1. The low mean similarity value of 0.0478 indicates minimal structural overlap with the training data, demonstrating that the model generates diverse linkers.

### 2.4. Fragment-Linker Attachment

The connection of a linker to an acyclic molecule is the final step in the MED framework for generating macrocycles. The acyclic molecule, symbolized by two cyclization sites marked by dummy atoms predicted in the transformer module, is connected with a linker derived from the EDM module.

#### 2.4.1 Attachment point validation and bond formation

The linking process begins with an acyclic molecule and the linker, each containing the required cyclization points, and converts them into molecular structures. If the input is invalid, it is discarded. To verify chemical plausibility, dummy atom positions are evaluated to ensure they are connected to atoms with available valence. They acquire it only when a designated attachment point is created (e.g., by replacing a hydrogen with a dummy atom). When a dummy atom is bound to an atom whose valence is already satisfied (e.g., oxygen), it is reassigned to an atom that has free bonding capacity, as shown in **Fig. 5**. This adjustment, combined with checks for chemical correctness, ensures attachment points allow new bonding without violating valence requirements.

**Fig. 5:**
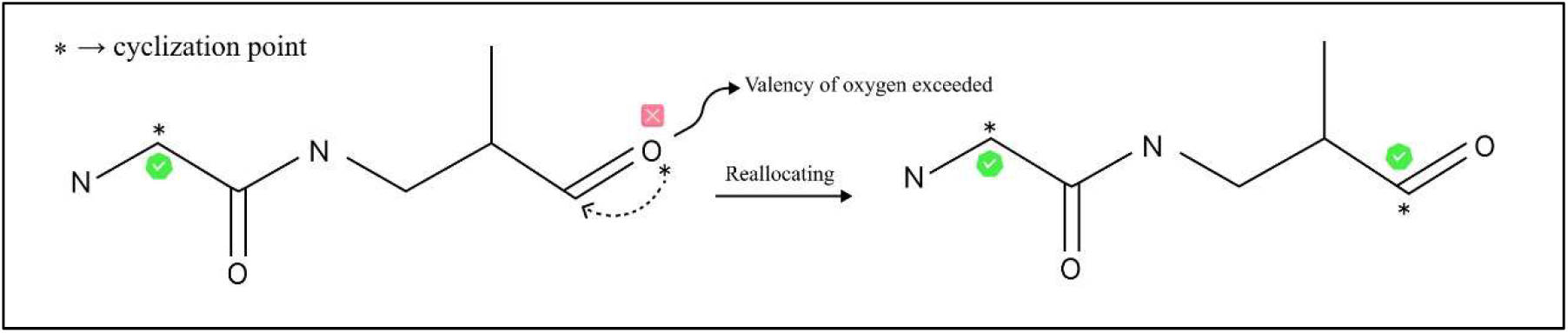
Reallocation of cyclization points. The illustration in Figure 5 shows how dummy atom (*) attachment points are moved from atoms with no bonding capacity left, such as from oxygen that has exceeded its valence, to a neighboring atom with available valence. This change is reflected in the image, ensuring the molecular structure stays chemically valid.

Both molecules contain dummy atoms, which must appear exactly twice each. Once these criteria are satisfied, the input molecules are merged into a single, disconnected structure within the molecular graph. If this condition is not fulfilled, the linking procedure is halted to ensure structural consistency and preserve the integrity of the molecular graph. The new single bond connections are then formed between the respective acyclic and the linkers’ dummy atoms that close the macrocyclic ring. After each bond is formed, dummy atoms are replaced and atomic indices are reset to ensure that anchors are correctly connected. The molecule is then checked to confirm proper valence, connectivity, and to detect any chemical faults. If the molecule fails these checks due to valence problems or invalid connectivity, it is rejected, and only valid macrocycles advance. Finally, the molecule is converted into a canonical SMILES string.

#### 2.4.2. Macrocycle filtering

To ensure that generated structures are macrocycles, each SMILES is filtered to have at least one ring that has a minimum of eleven atoms, and any non-macrocyclic structures that may result from invalid linking are eliminated. Multiple linkers that are generated for each acyclic fragment are evaluated in fragment–linker pairs, enabling the exploration of diverse combinations. Duplicate macrocycles are eliminated by using unique SMILES and preserving them for scoring, which allows the model to become more precise and ensures that only chemically plausible macrocycles are generated. This attachment approach strikes a balance between flexibility and accuracy, yielding a predictable collection of macrocycles suitable for drug discovery^7^. A summary of the whole pipeline is depicted in **Fig. 6**.

**Fig. 6:**
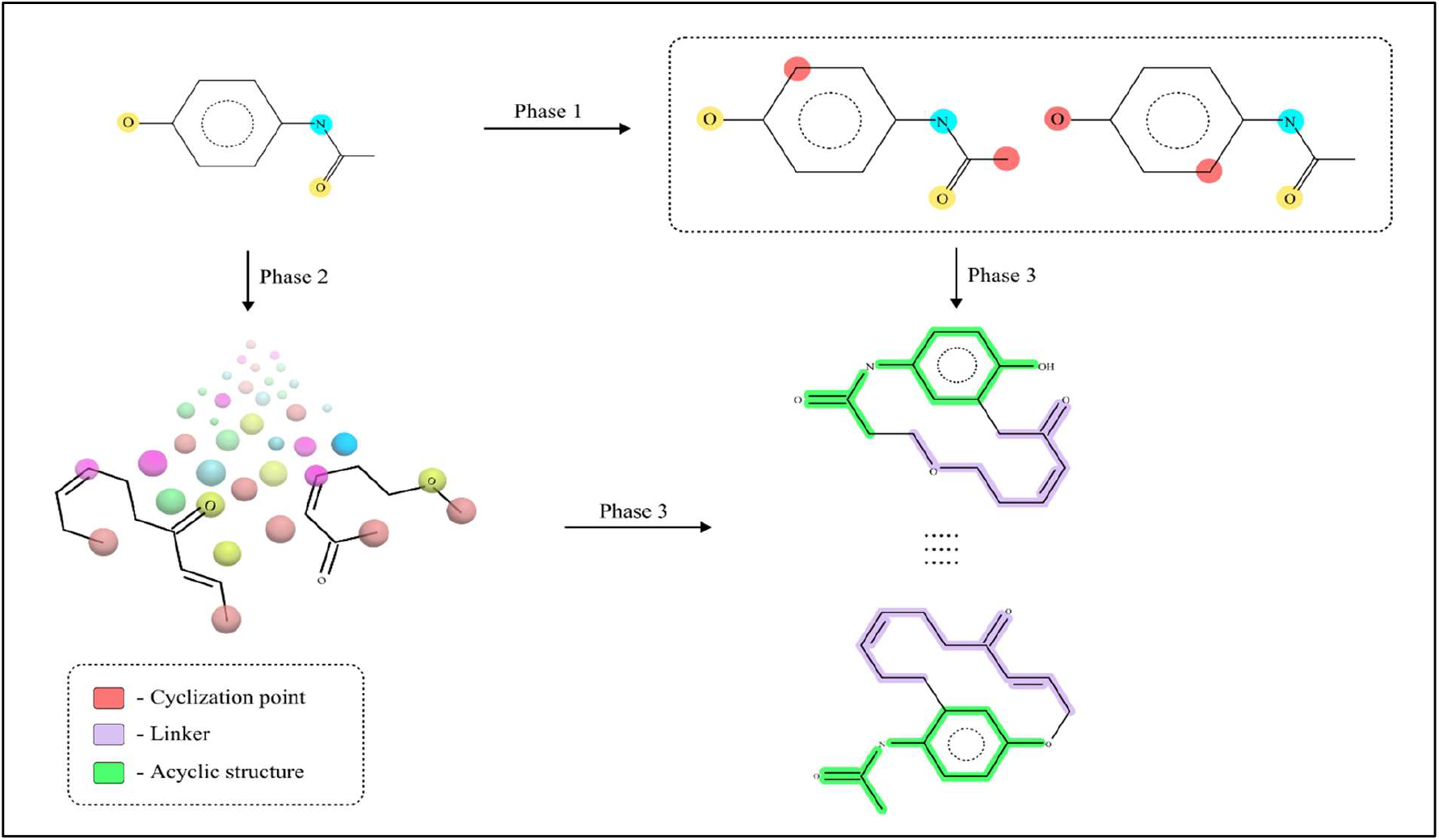
Computational pipeline for macrocyclization. In Phase 1, transformer-based prediction identifies cyclization sites on the acyclic structure. Phase 2 employs an E(3)-equivariant diffusion model to generate chemically consistent linkers, and in Phase 3, linkers are attached to form diverse macrocycles.

### 2.5. Docking, Screening, and Visualization of Protein–Ligand Complexes

As a case study, macrocyclic derivatives of the Janus Kinase-2 (JAK2) inhibitors^19,21,33^ Fedratinib, Ruxolitinib (Direct inhibitors), Sunitinib, and Lenvatinib (Indirect inhibitors) were designed and docked against the AlphaFold^34^-predicted JAK2 structure (Uniprot ID: O60674), which provides a complete and native-like protein conformation free from crystal-packing artifacts which are present in the crystal structure available in the Uniprot, having an amino acid length of 1132. The JAK2 structure selected has per-residue confidence scores (average pLDDT^35^: 87.22), ensuring structural reliability for binding site analysis. Standard protein preparation protocols were followed, including removal of non-essential molecules, addition of hydrogens, charge assignment, and energy minimization. Docking simulations were performed on macrocyclic structures generated for acyclic JAK2 inhibitors using PyRx^36^ with the AutoDock Vina algorithm, employing grid boxes centered on the ATP-binding pocket to explore potential binding orientations. The best-scoring ligands were subsequently analyzed in Biovia Discovery Studio 2025^37^ to map key non-covalent interactions, including hydrogen bonding, hydrophobic contacts, and π–π stacking^38^, with JAK2 residues. Conformational analyses assessed macrocycle stability and binding site adaptability, providing insights into binding behavior.

## 3. Results and Discussion

### 3.1. Performance Metrics

MED was tested with a reduced two-stage setup, where transformer-based site prediction was directly followed by equivariant diffusion generation, bypassing the fragment-linker attachment step entirely. Without explicit attachment rules governing how fragments and linkers connect, the model often failed to close rings correctly or generate stable macrocyclic structures. This configuration achieved a chemical validity of 98.74% and a uniqueness score of 91.67%, but a limited macrocyclization percentage of 35.73%.

By contrast, the full three-stage MED pipeline overcame these limitations: the fragment–linker attachment module proved essential for chemically valid closures and for retaining drug-like macrocycles.

**Table 1.** summarizes the comparative performance of Macformer, MacLS, and MED on macrocyclization tasks, evaluated on 5,551 samples of the ZINC dataset. Metrics for Macformer and MacLS are taken directly from Diao et al. (2023)^10^. Metrics include validity, uniqueness (among valid molecules), macrocyclization success (among valid and unique molecules), and novelty of predicted linkers, reflecting structural accuracy and chemical plausibility.

**Table 1.** Comparison of metrics with various models.

| Metrics | MED | Macformer | MacLS |
| --- | --- | --- | --- |
| Validity | 93.82±1.02% | 72.91±2.05% | 89.67±0.07% |
| Uniqueness | 99.94±0.02% | 47.74±8.98% | 95.04±0.14% |
| Macrocyclization % | 99.92±0.05% | 96.39±0.71% | 100±0.00% |
| Novelty of the linker | 82.81±2.12% | 44.24±2.05% | 0.00±0.00% |

To further assess the chemical plausibility of the transformer-based attachment point predictions, the standalone transformer model was tested separately on the ZINC database^9^. It achieved a validity of 99.3175%, a uniqueness score of 99.9595%, and a novelty of 96.7684%. Additionally, the average Tanimoto similarity between predicted and input molecules was 0.9119, indicating a high degree of structural preservation while introducing the desired cyclization. These results confirm that the transformer outputs are chemically valid, providing a strong foundation for the overall MED pipeline.

#### Model evaluation metrics

##### Validity

The percentage of generated chemically valid molecules.

##### Uniqueness

The percentage of unique molecules in the generated valid molecules.

##### Novelty of linker

The percentage of novel linkers that are not present in the training set, in the generated valid and unique molecules.

##### Macrocyclization percentage

The percentage of macrocycles in the generated valid and unique molecules.

### 3.2. Feature Distributions of Generated Molecules

To evaluate the structural validity, diversity, and physicochemical realism of the molecules generated by MED, We evaluated the model on 1,000 molecules sampled from the ZINC dataset across a selected set of molecular descriptors. The corresponding feature distributions are presented in the Supplementary Figures (Fig. SF2 Fig. SF8).

#### 1. Normalized Principal Ratios (NPR1, NPR2)

NPR1 and NPR2^39,40^ are used to help quantify molecular shapes by distinguishing between rod-like, disc-like, and spherical geometries as shown in **Supplementary Fig. SF2**. NPR descriptors enable comparison of shape diversity and ensure that generated molecules cover a broad region of shape space rather than collapsing to a single geometry.

#### 2. Plane of Best Fit

The PBF^41^descriptor measures molecular planarity, which is relevant for π-stacking interactions, aromaticity, and conformational preferences. Including PBF (**Supplementary Fig. SF3**) allows evaluation of whether the model preserves realistic 3D characteristics rather than generating overly planar or overly distorted structures.

#### 3. Bond lengths

Bond length distributions serve as a structural validity check by ensuring that generated molecules lie in chemically reasonable interatomic distances^8^. The values as shown in **Supplementary Fig. SF4** remain within chemically reasonable ranges, indicating that the model produces physically plausible linker geometries.

#### 4. Energy per atom

Energy per atom provides a normalized measure of molecular stability. The energy per atom distribution (**Supplementary Fig. SF5**) shows that the generated molecules generally have reasonable stability, indicating that the model does not produce energetically unfavorable structures.

#### 5. Number of Torsions

The number of torsions or the number of rotatable bonds represents how flexible a molecule is. Higher the values greater the conformational freedom. The distribution of torsion counts for the generated molecules is shown in **Supplementary Fig. SF6** reflecting that the model produces molecules with varying levels of flexibility.

#### 6. LabuteASA

Labute Accessible Surface Area (LabuteASA)^42^ represents the solvent-accessible surface area of molecules reflecting its overall size and exposure to the surrounding environment as shown in **Supplementary Fig. SF7**. Higher values indicate increased structural diversity within drug-like chemical space.

#### 7. Radius of gyration (Rg)

The Rg captures the overall spatial extent and compactness of a molecule (**Supplementary Fig. SF8**). Rg is useful for assessing molecular size and conformational spread, which are important for comparing generated molecules with the training ones^43^.

### 3.3. Docking, screening, and visualization studies with the JAK2 protein

As a focused case study, docking and pharmacokinetic screening were carried out on selected MED-generated macrocycles derived from known JAK2 inhibitors to evaluate their structural validity, and suitability for practical synthesis. Docking studies against the high-confidence AlphaFold-predicted JAK2 structure identified multiple macrocyclic derivatives (F1, S1, R1, and L1), especially the top structures with good pharmacokinetic properties derived from Fedratinib (F), Ruxolitinib (R), Sunitinib (S), and Lenvatinib (L). These derivatives showed predicted binding affinities in the highly favorable range (-7.8 to -10.6 k.cal. mol^−1^), specifically binding to the ATP binding pocket (Lys882), the proton acceptor site (Asp976), and the region from Leu855 to Val863 as specified in Uniprot. Improved binding within the generated grid (center in Å: X: 28.09, Y: 9.50, Z: -33.37; dimensions in Å: X: 32.29, Y: 16.74, Z: 19.69) was consistently linked to optimized hydrogen bonding to hinge residues, increased hydrophobic interactions within the ATP-binding pocket, and conformational preorganization that reduced entropic penalties upon binding. Structural visualization using Biovia Discovery Studio 2025 confirmed better binding pocket occupancy for several macrocyclic analogues compared with acyclic precursors, indicating potential for enhanced potency and selectivity.

Table 2 presents the 2D and 3D interactions between JAK2 interacting amino acid residues with the structures generated and visualized by Biovia Discovery Studio 2025. Where J_F: JAK2_Fedratinib complex; J_F1: JAK2_Macrocyclic Fedratinib complex; J_S: JAK2_Sunitinib complex; J_S1: JAK2_Macrocyclic Sunitinib complex; J_R: JAK2_Ruxolitinib complex; J_R1: JAK2_Macrocyclic Ruxolitinib complex; J_L: JAK2_Lenvatinib complex; J_L1: JAK2_Macrocyclic Lenvatinib complex; J_P(DB): JAK2_Pacritinib (PMID: 35983661) (PMID: 35993369) (Drugbank) complex; J_P(MF): JAK2_Pacritinib (MacFormer) complex.

**Table 2.**
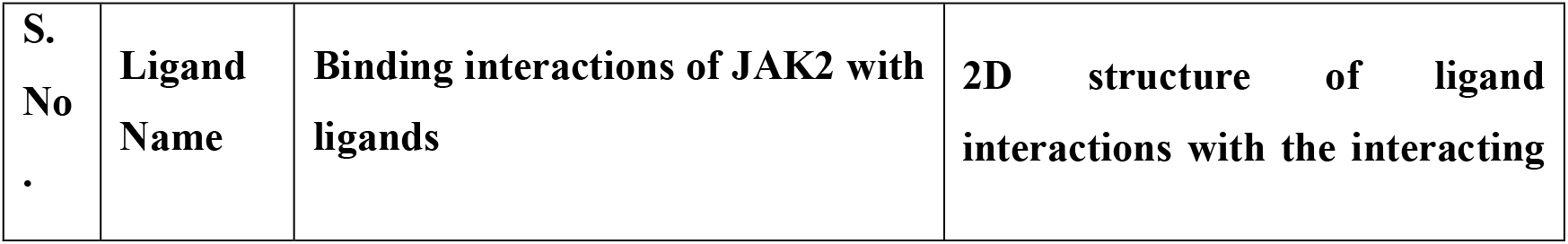
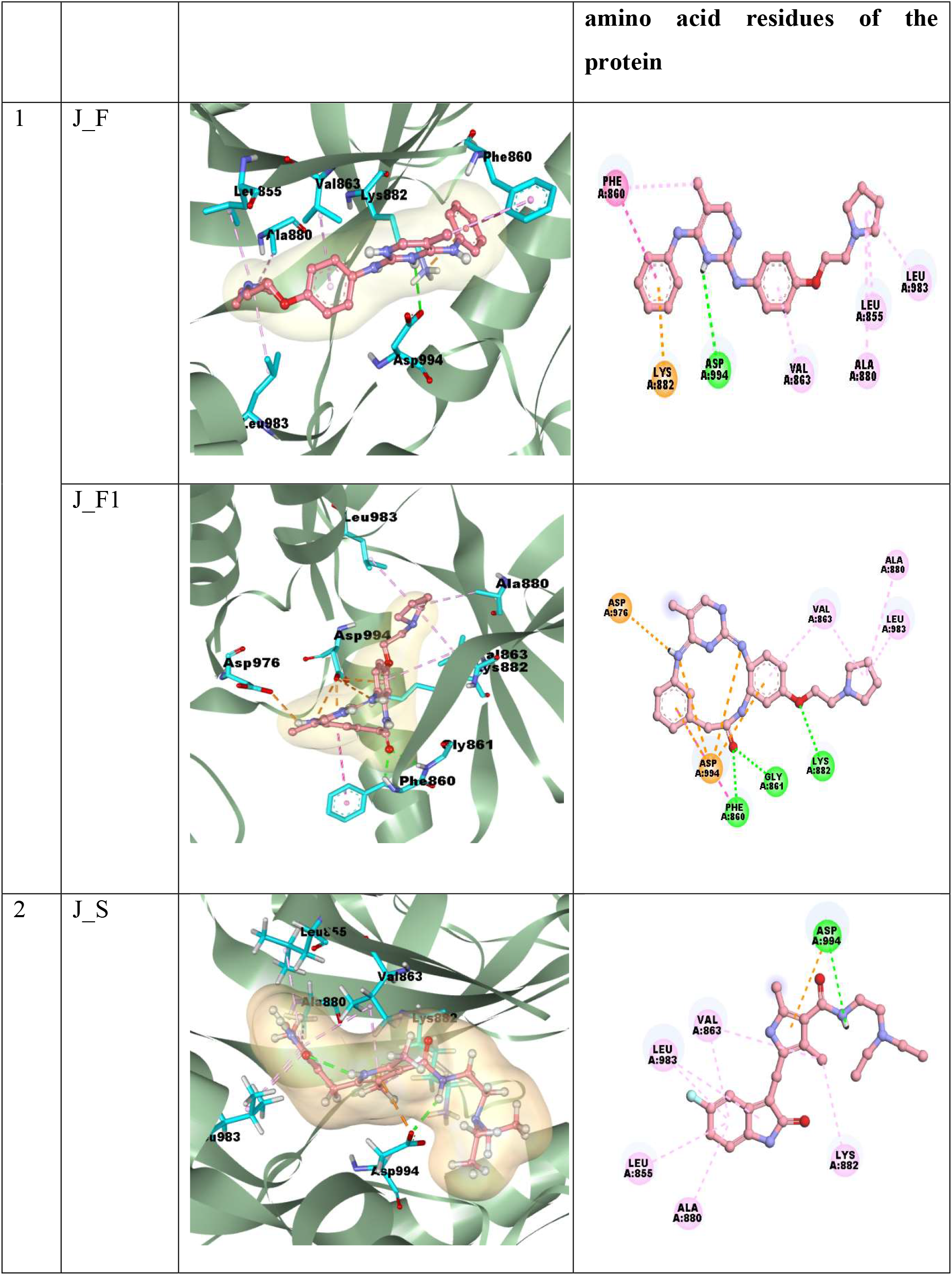
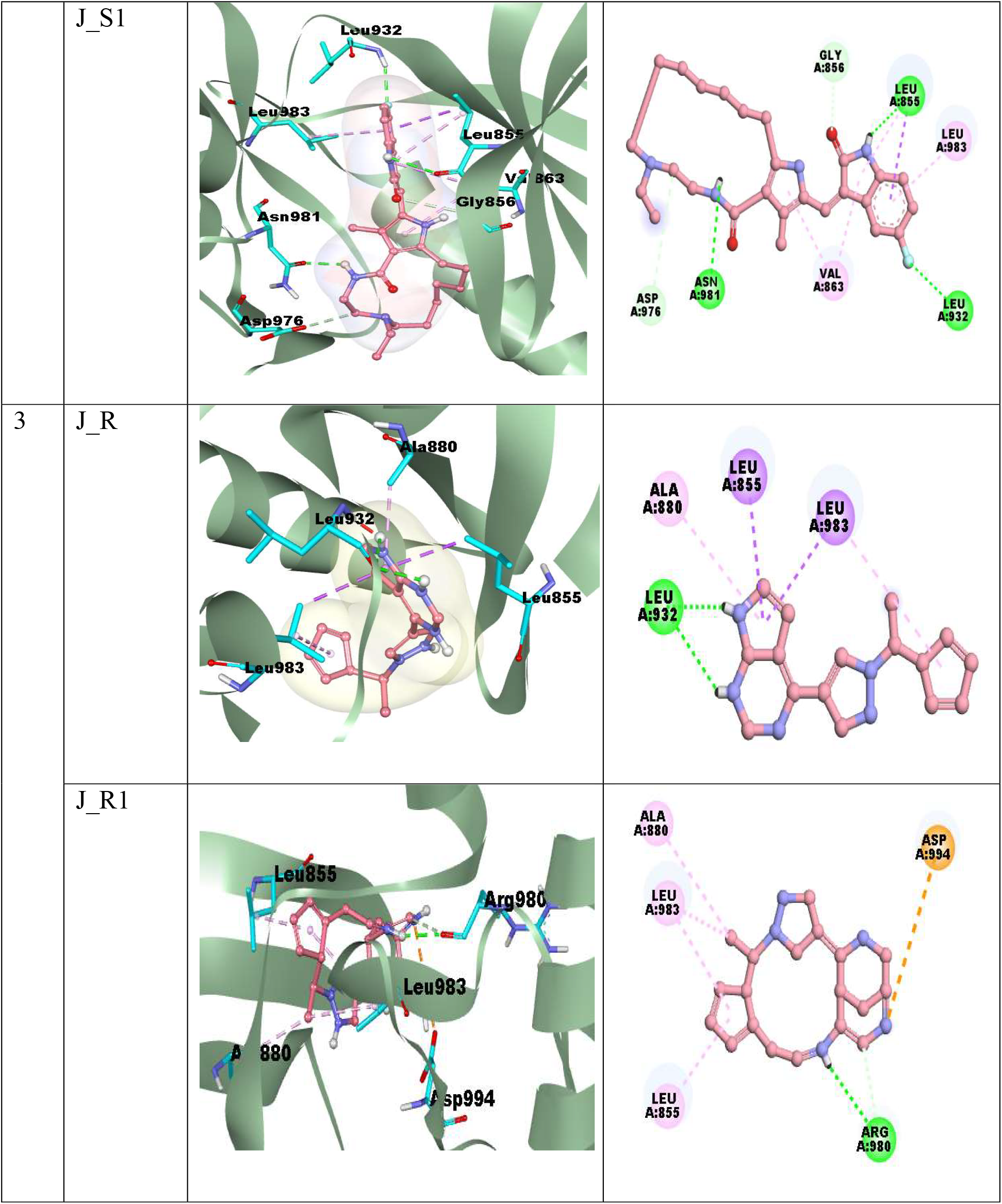
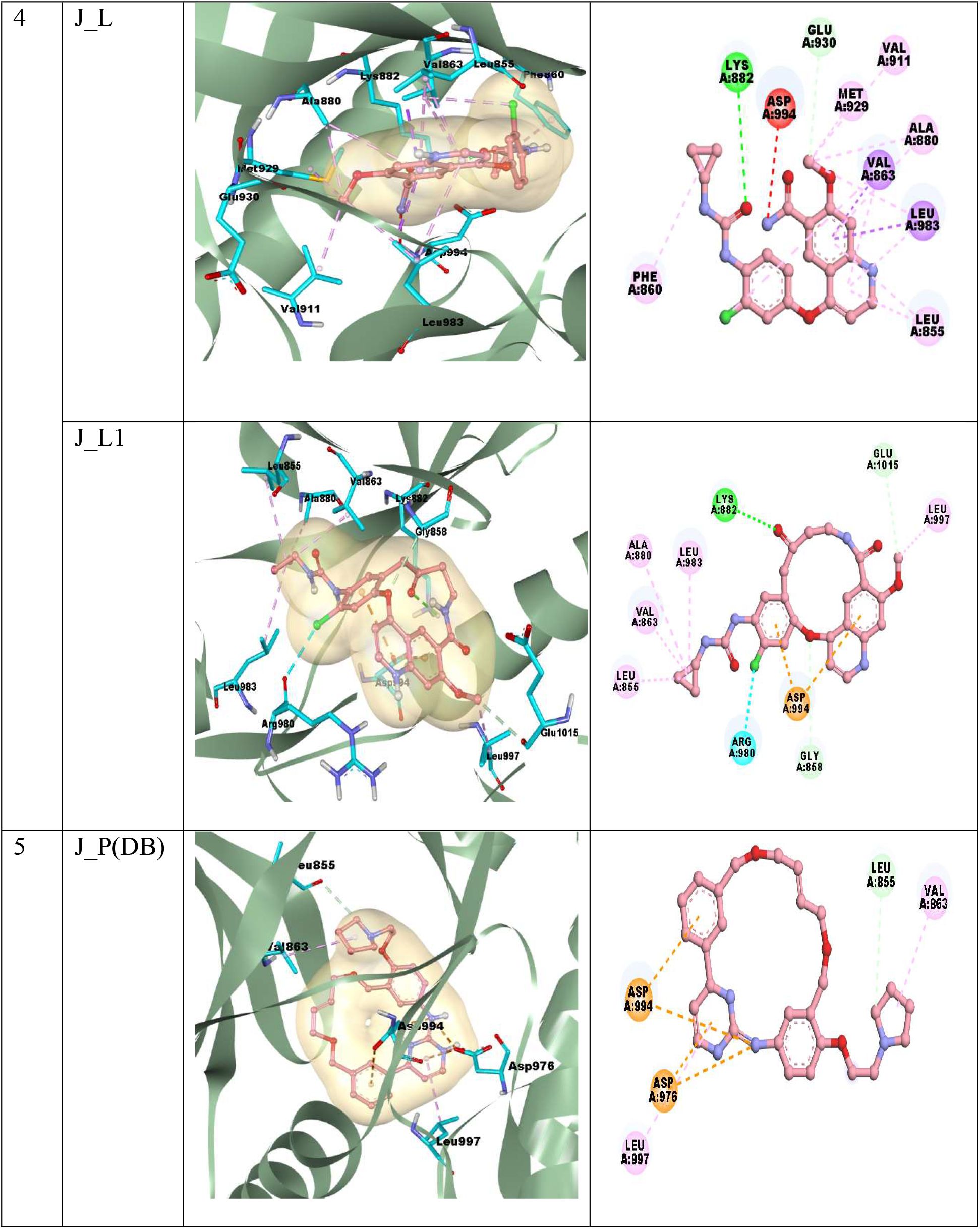
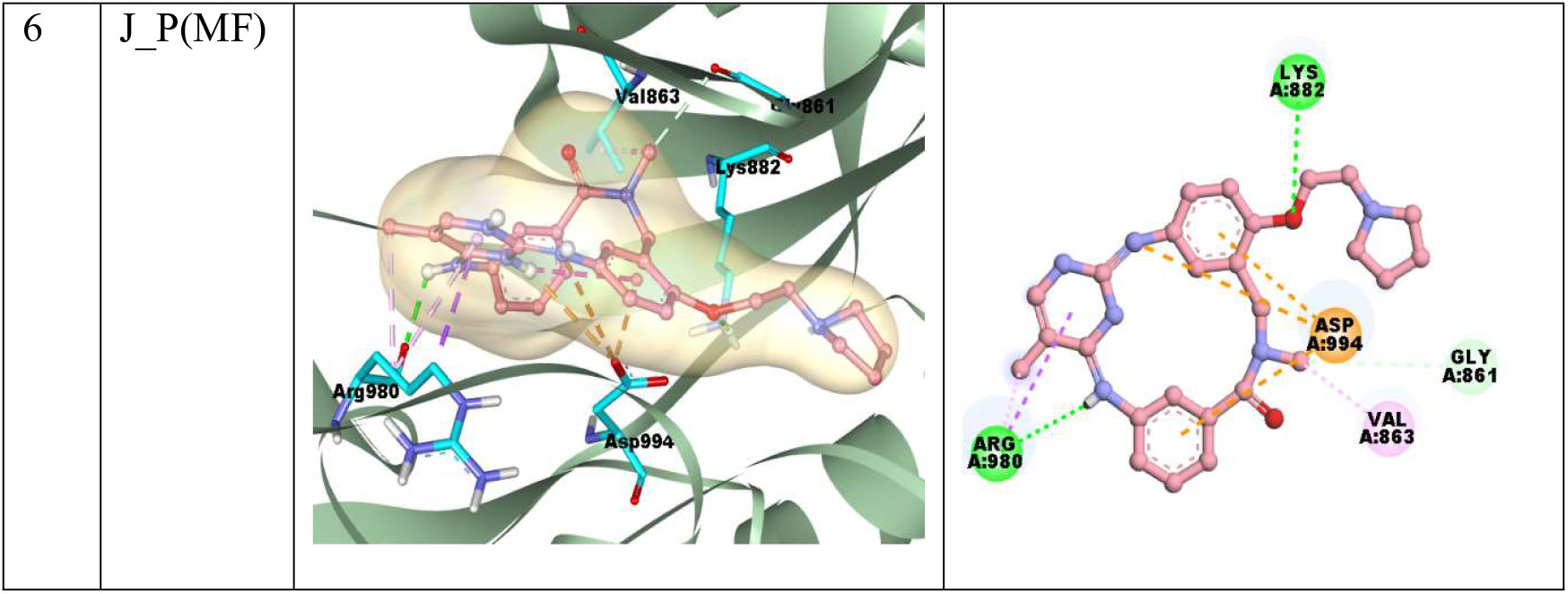
Binding interactions of JAK2 interacting amino acid residues with the ligands.

Table 3 summarizes the binding affinity scores (obtained from PyRx), the structures of ligands, and the interactions between the JAK2 protein and ligands in angstroms (Å). Where F: Fedratinib,F1: Macrocyclic Fedratinib, S: Sunitinib, S1: Macrocyclic Sunitinib, R: Ruxolitinib, R1: Macrocyclic Ruxolitinib, L: Lenvatinib, L1: Macrocyclic Lenvatinib, P(DB): Pacritinib from Drugbank, P(MF): Pacritinib generated through Macformer. Structures and interactions are generated by Biovia Discovery Studio 2025.

**Table 3.**
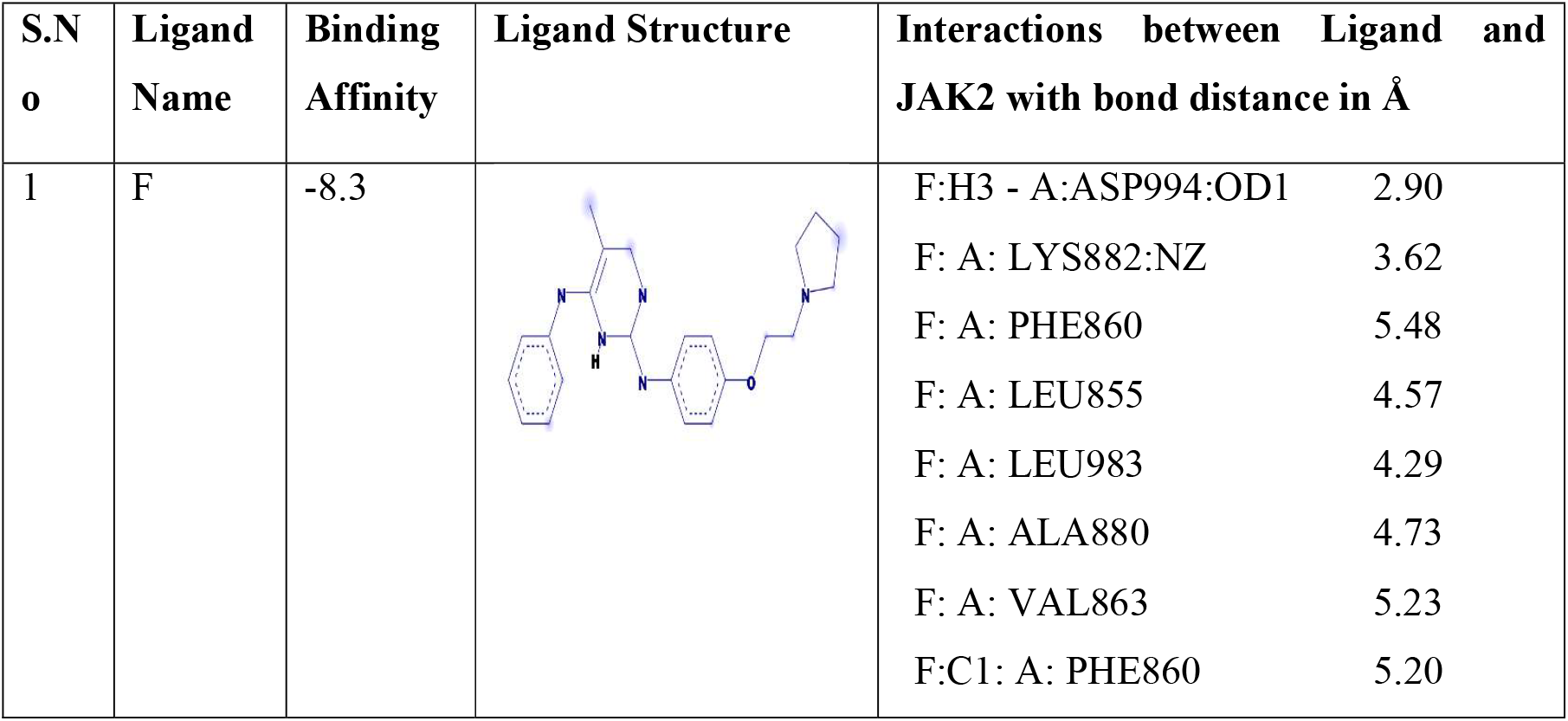
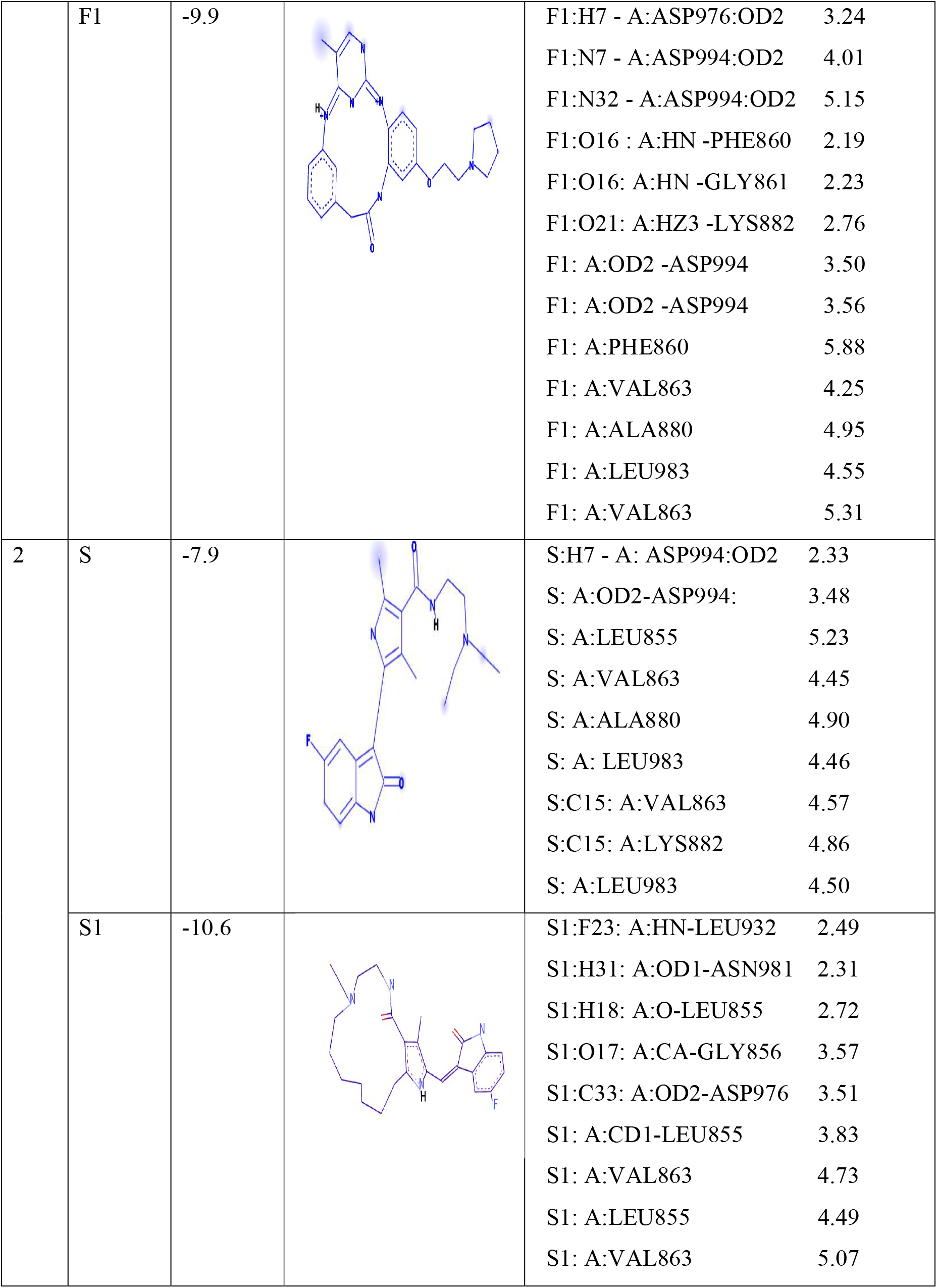
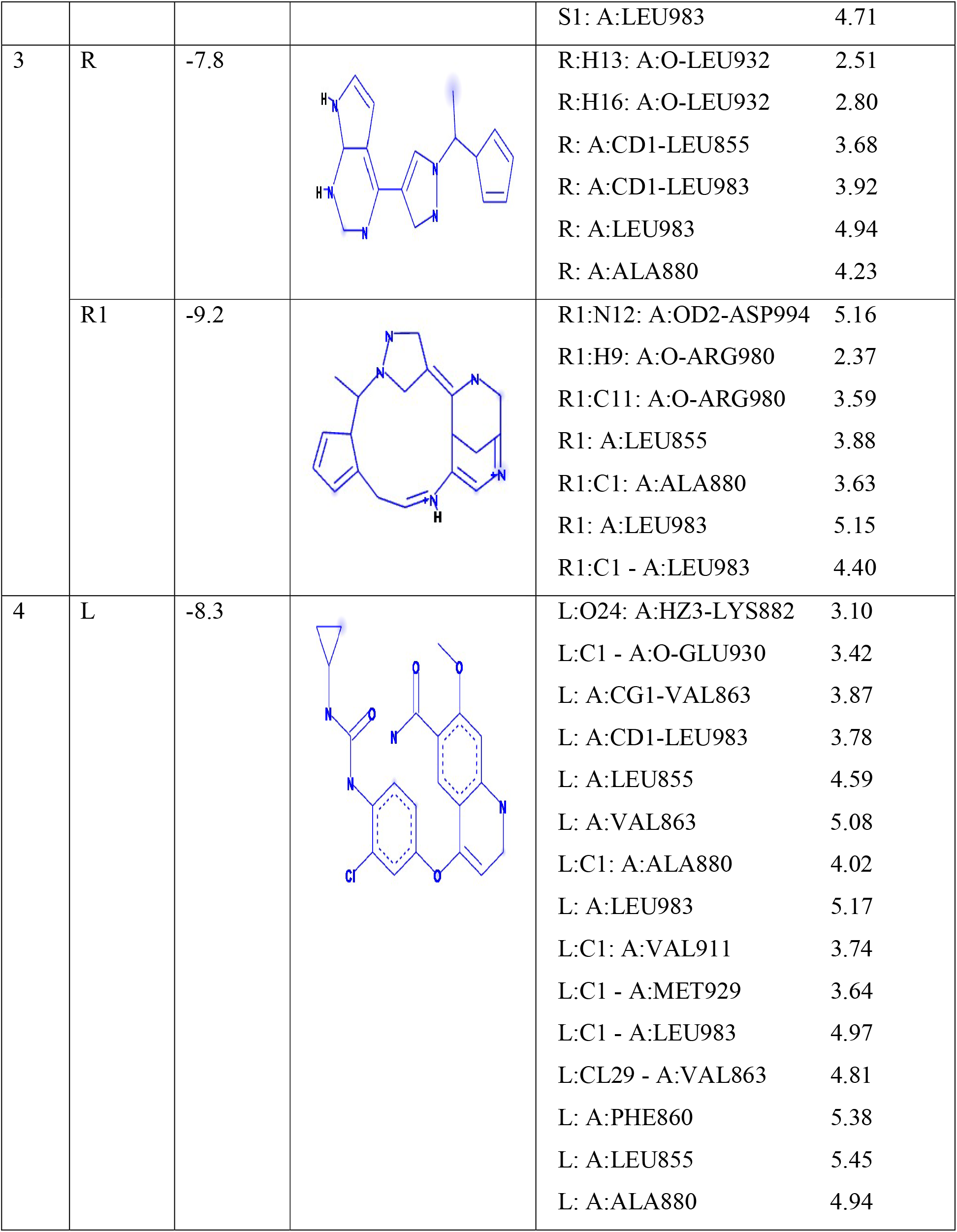
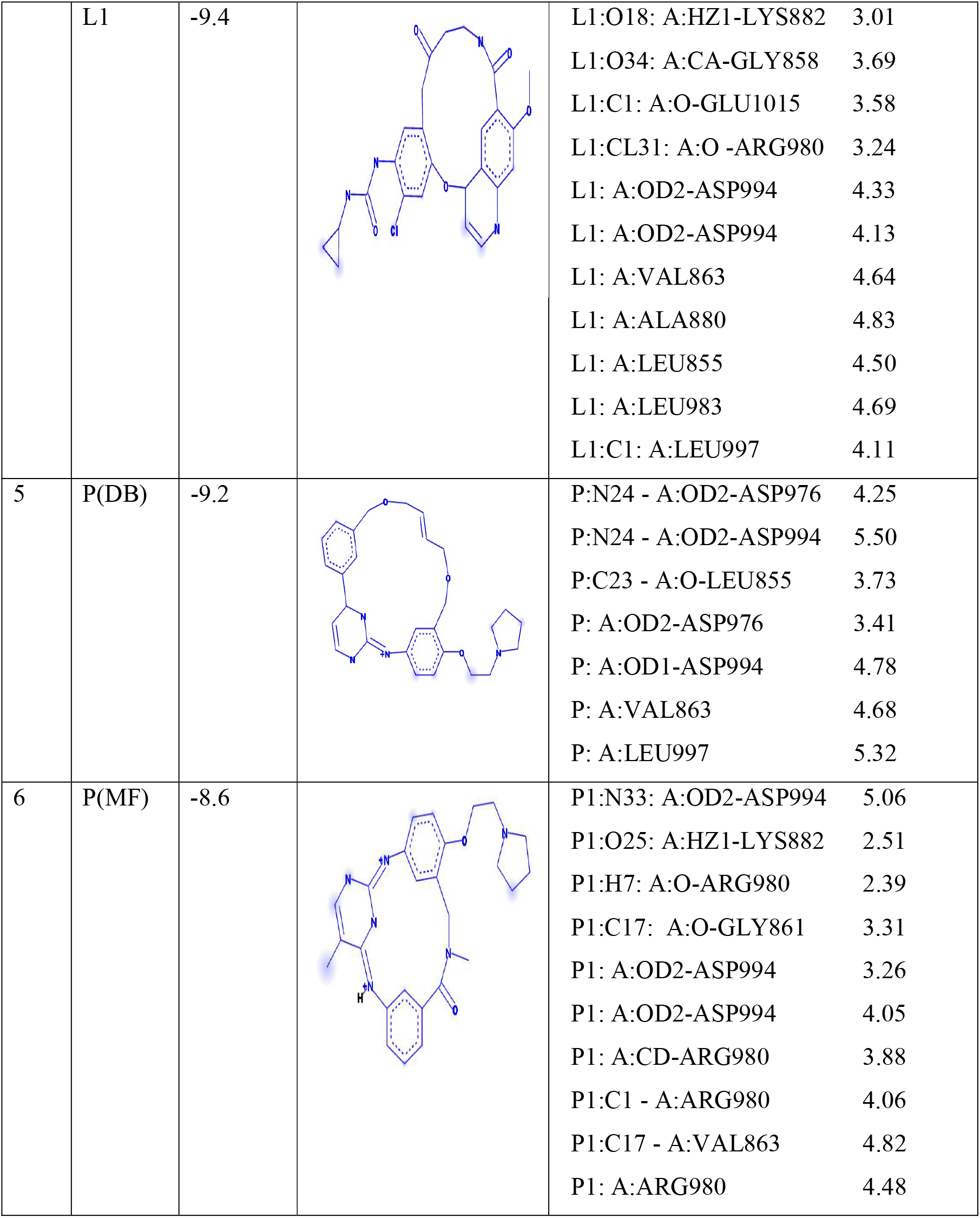
Interactions between ligands and the JAK2 protein.

Table 4 summarizes the pharmacokinetic properties of the generated structures and the cyclic drugs produced by ADMETlab 2.0 (PMID: 38572755). The valid ranges for these properties are as follows: MW (Molecular weight in grams per mole): less than 750; nHA (number of Hydrogen bond Acceptors): ≤10; nHD (number of Hydrogen bond Donors): ≤5; TPSA (Total Polar Surface Area in Å^2^): 7 to 200; logP: ≤5.0; caco2 (cell permeability in gut-blood barrier): 5 to less than -5.0 cm/sec; BBB (Blood Brain Barrier): -3.0 to 1.2; QED (Quantitative Estimate of Drug-likeness): 0 to 1; logS (aqueous solubility): -6.5 to 0.5; logD (distribution coefficient): 1 to 3 (hydrophilic), 3 to 5 (lipophilic).

**Table 4.**
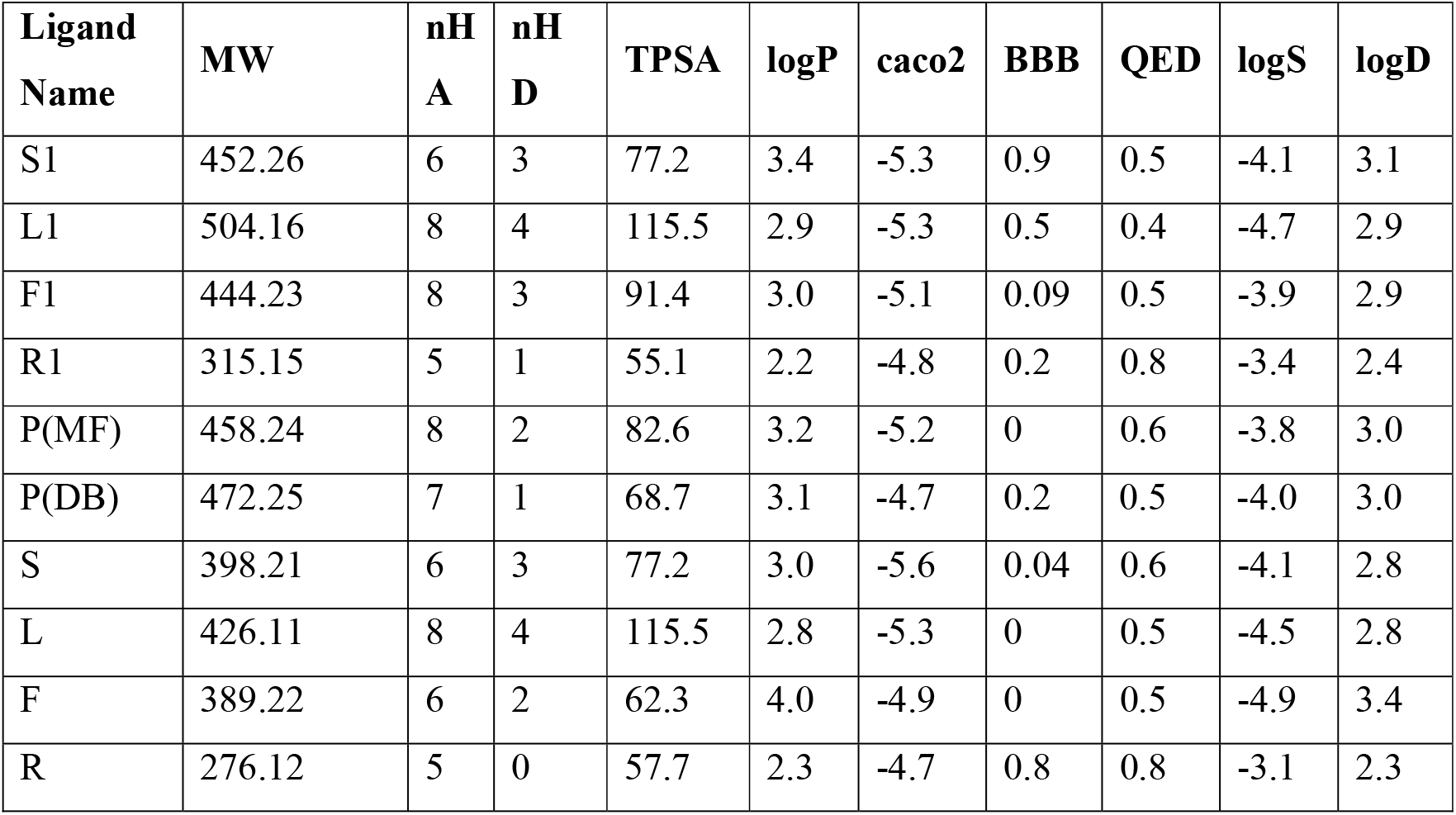
Pharmacokinetic Properties of the generated Macrocyclic structures and acyclic drugs.

Fedratinib uniquely inhibits the thiamine transporter SLC19A3^44^ by structurally mimicking thiamine. Its 2,4-diaminopyrimidine core, sulfonamide branch, terminal pyrrolidine, and hydrophobic tert-butyl group collectively align with thiamine’s binding determinants, allowing it to occupy the transporter pocket competitively. This off-target interaction explains the observed reduction in thiamine uptake and the associated risk of Wernicke’s encephalopathy^45^ in patients treated with this drug. In contrast, the macrocyclic structures of Ruxolitinib (R1), Sunitinib (S1), and Lenvatinib (L1) are based on chemically distinct scaffolds and lack the pharmacophoric elements needed to mimic thiamine. Consistent with their structures, these predicted macrocyclic structures do not significantly inhibit SLC19A3, making them more selective for their intended kinase targets. Therefore, in the context of JAK2 inhibition, alternative options with macrocyclic structures (S1, R1, and L1) may be preferred over Fedratinib to lower the risk of transporter-mediated thiamine deficiency. Overall, these findings demonstrate that model-guided macrocyclization, combined with structure-based design, enables the rapid creation of synthetically accessible scaffolds with enhanced binding properties. The improved binding observed, driven by preorganization and optimized interaction networks, offers strong support for applying this strategy to other therapeutic targets. This proof-of-concept workflow demonstrates the potential of AI-driven molecular design to accelerate the discovery of next-generation macrocyclic inhibitors.

## 4. Conclusion

In this work, we present Macro-Equi-Diff (MED), a hybrid transformer-diffusion framework for macrocyclization of drug-like molecules from their acyclic precursors. MED integrates a transformer-based module for predicting cyclization sites, an EDM for linker generation, and a fragment–linker attachment module. This guarantees that the generated macrocyclic structures are chemically valid and geometrically consistent. Our results demonstrate that MED surpasses existing methods in terms of validity, uniqueness, novelty, and macrocyclization, while also generating macrocycles with enhanced drug-like properties.

To further evaluate the structural realism and diversity of the generated molecules, we analyzed multiple geometric and physicochemical feature distributions, including NPR1, NPR2, PBF, Bond lengths, energy per atom, number of torsions, LabuteASA, and radius of gyration. A case study on the JAK2 protein demonstrated how MED can generate macrocycles, yielding diverse analogs with favorable docking results. Importantly, MED’s primary goal is to generate valid macrocycles; further research will be necessary to assess the therapeutic potential of these macrocycles fully.

Overall, MED provides chemists with a structured approach to explore macrocyclic chemical space, starting with acyclic scaffolds and progressing towards synthetically accessible macrocycles. With geometric insight and diffusion-based modeling enhancing the process, MED provides an adaptable framework that could speed up the discovery of next-generation macrocyclic molecules.

## Abbreviations

ADMET: Absorption, Distribution, Metabolism, Excretion, and Toxicity
ATP: Adenosine Triphosphate
COM: Center of Mass
CYP: Cytochrome P4504
DS2025: Biovia Discover Studio 2025
E(3): Euclidean Group in 3 Dimensions (translations, rotations, reflections)
EGCL: Equivariant Graph Convolutional Layer
EGNN: Equivariant Graph Neural Network
GNN: Graph Neural Network
HBA: Hydrogen Bond Acceptor
HBD: Hydrogen Bond Donor
hREG: Human Ether-à-go-go-Related Gene
JAK2: Janus kinase 2
MacLS: Macrocycle Library Screening
MED: Macro Equi Diff
MLP: Multi-Layer Perceptron
MW: Molecular Weight
O(3): Orthogonal group in three dimensions, representing 3D rotations and reflections.
PDBQT: Protein Data Bank, Quaternion, Torsion
pLDDT: predicted Local Distance Difference Test
PSA: Polar Surface Area
PyRx: Python Prescription
QED: Quantitative Estimate of Drug-likeness
Ro5: Rule of 5 (drug-likeness guideline in medicinal chemistry)
SA: Synthetic Accessibility
SDF: Structure Data File
SMILES: Simplified Molecular Input Line Entry System”

## ACKNOWLEDGEMENTS

We authors, S.S.K, S.A, S.L.G, V.R.K, S.G, and V.K, express our sincere gratitude to the Drugparadigm Research Lab for providing the necessary facilities and infrastructure that enabled the successful completion of this work.”

## AUTHOR CONTRIBUTIONS

S.S.K., S.A., S.L.G., and V.R.K. contribute equally to the work. S.S.K. and V.R.K. performed investigation, validation, methodology, visualization, writing—original draft, and formal analysis; S.A. and S.L.G. conducted data curation, formal analysis, methodology, validation, and visualization; S.G. worked on conceptualization, methodology, validation, visualization, formal analysis, data curation, software, supervision, writing—review, and editing; V.K. analyzed the *in silico* studies methodology, including writing review, editing, and supervision; all the authors edited and approved the manuscript.

## ADDITIONAL INFORMATION

Correspondence should be addressed to Vani Kondaparthi.

## AVAILABILITY OF DATA

All the reported data are available in the links and references provided in the manuscript.

## CODE AVAILABILITY

The code is available at https://github.com/drugparadigm/Macro-EquiDiff

