## Supplementary Information for "Macro-Equi-Diff (MED): Scaffold-based Macrocycles Generation Using Equivariant Diffusion"

1. **Algorithm: Macrocyclization Workflow in the MED Model**

2. Input:
   a. Acyclic molecule SMILES string: s
   b. Hyperparameters: Number of augmented variants per input molecule ($N_{AUG}$ ), linker size, minimum ring size($min_{ring}$) = 11
3. Output:
   a. Top macrocycles ranked by drug-likeness score, with biopharmaceutical properties.
   b. Intermediate files: Augmented SMILES, cyclized fragments, linkers, and final macrocycles.
4. Phase I: Preprocessing and Augmentation
   a. Neutralize radicals: s → $s_{clean}$
   b. Augment SMILES: $s_{clean}$ → A = ($a_1, . . . , a_{N_{AUG}}$)
5. Phase II: Cyclization Point Prediction (Transformer Module)
   a. Predict two anchor points: ∀$a_i$ ∈ A → $c_i$ ∈ C
6. Phase III: Linker Generation (Diffusion-Based Module)
   a. Parameters: Linker size, pre-trained checkpoint.
   b. Generating linker:
      i. Graph representation: Nodes (atoms) + edges (bonds) → 3D coordinates.
      ii. Embedding: Node/edge features
      iii. EGNN layers:Edge update = MLP on concatenated features and Node update = aggregate edges + MLP.
      iv. Outputs: ∀$a_i$ ∈ A → L = ($l_1$, . . . )
7. Phase IV: Fragment-Linker Attachment
   a. Connect: For each $c_i$ ∈ C, bonds with $l_j$ ∈ L are validated
8. Phase V: Property Calculation
   a. Molecular Descriptor Calculation and Evaluation of Validity, Uniqueness, and Novelty

### 2. Tanimoto similarity

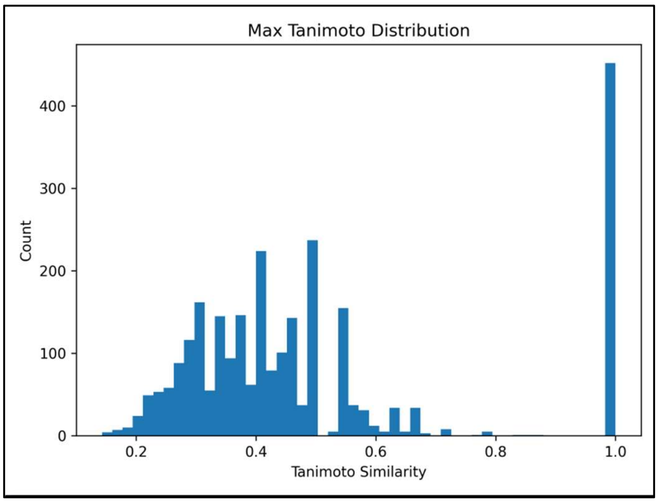


**Supplementary Fig. SF1:** Average Tanimoto similarity between generated linkers and the training set.

The mean Tanimoto similarity across generated linkers is 0.0478, indicating low global similarity to the training distribution. This behavior indicates that the model does not collapse to a narrow region of chemical space. Instead, the results indicate that the model learns linker chemistry while generating diverse and novel molecules without overfitting to the training data.

**3. Feature distributions of generated molecules.**

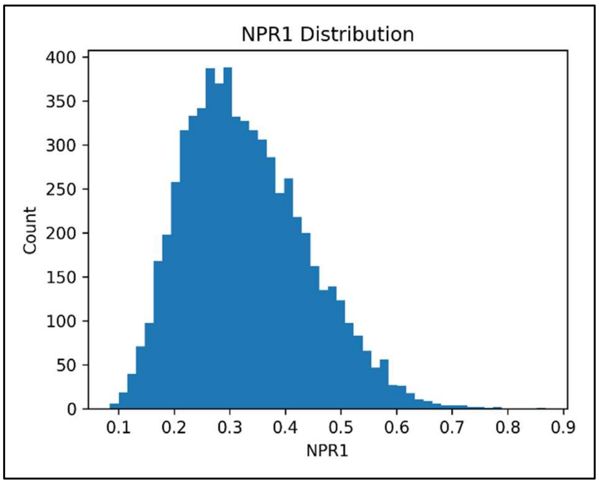


**Supplementary Fig. SF2.a.** NPR1 distributions of generated molecules.

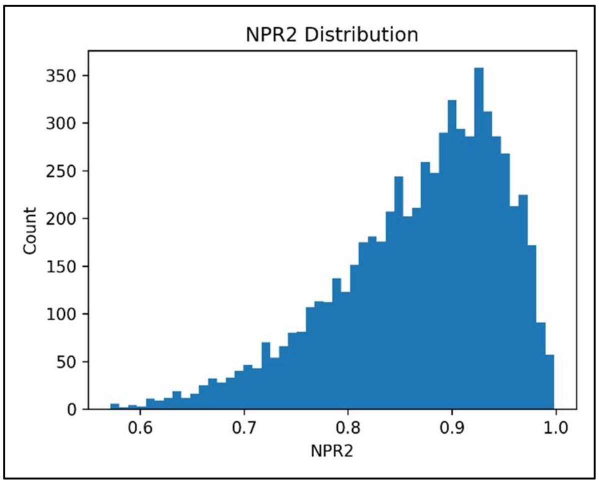


**Supplementary Fig. SF2.b.** NPR2 distributions of generated molecules.

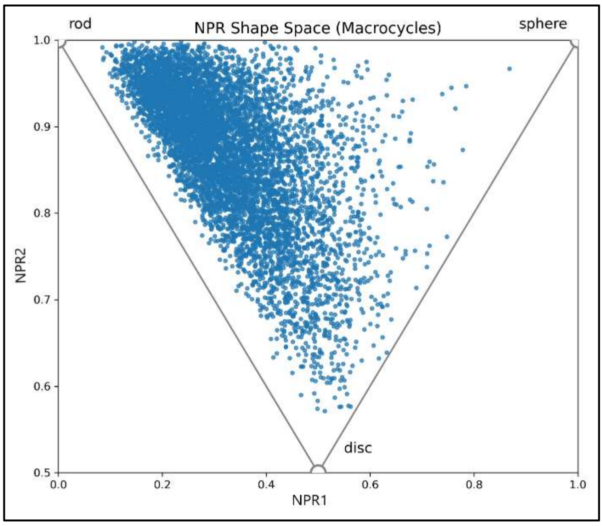


**Supplementary Fig. SF2.c.** NPR1 vs NPR2.

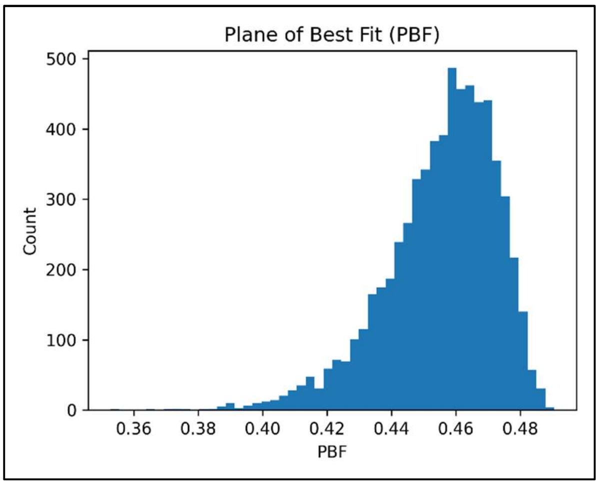


**Supplementary Fig. SF3.** Plane of Best Fit (PBF) values illustrating molecular planarity.

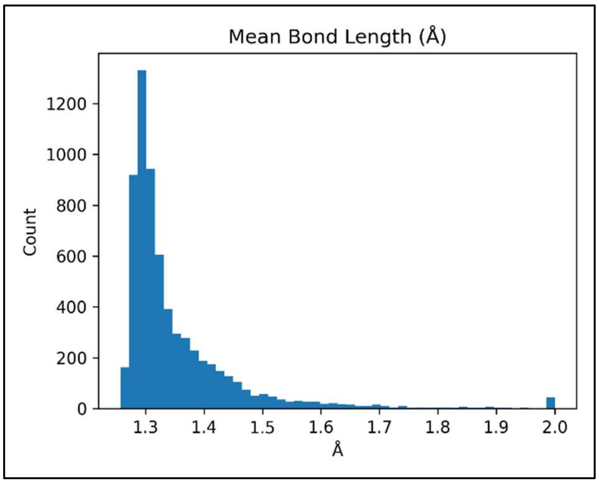


**Supplementary Fig. SF4.** Mean bond length values (in Armstrongs) across generated molecules.

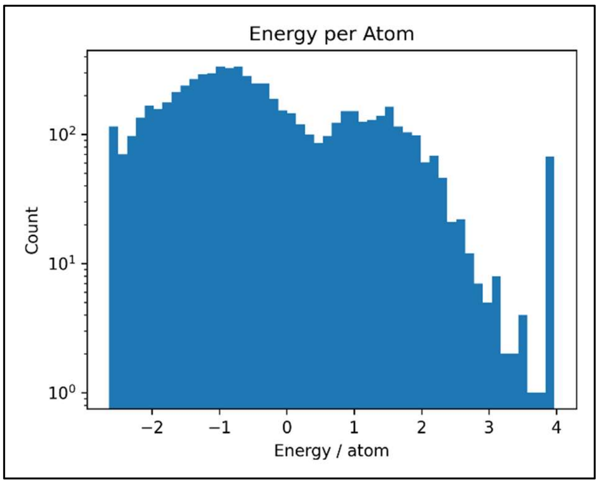


**Supplementary Fig. SF5.** Energy per atom distribution of generated molecules.

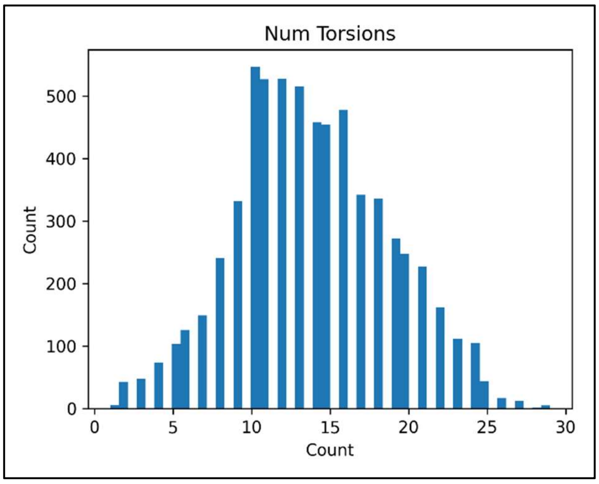


**Supplementary Fig. SF6.** Distribution of the number of torsions in generated molecules.

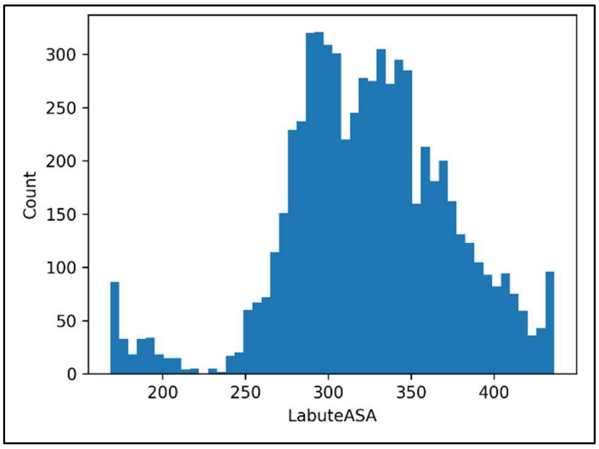


**Supplementary Fig. SF7.** Labute Approximate Surface Area (LabuteASA) distribution.

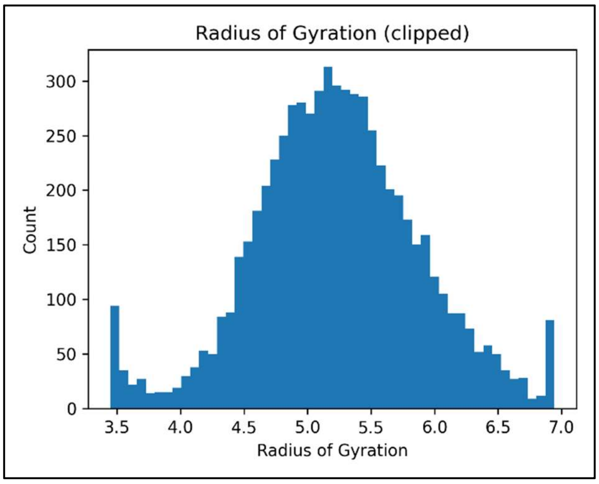


**Supplementary Fig. SF8.** Radius of gyration distribution of generated molecules.
